# Efficient long-term cryopreservation of *Arthrospira platensis* NIES-39 and adaptation for the Space Algae-2 spaceflight experiment

**DOI:** 10.1101/2025.09.02.671390

**Authors:** Katherine E. Fisher, A. Mark Settles

## Abstract

*Arthrospira platensis*, commonly known as spirulina, is a photosynthetically efficient cyanobacterium used for production of nutritional supplements, biofuel, wastewater treatment, and CO_2_ sequestration. These traits and uses make it a candidate for bioregenerative life support systems for space travel and future habitats. However, *A. platensis* lacks a cryopreservation method for long-term storage of viable cells, and genetic stocks are currently sub-cultured for maintenance. Here we report an efficient cryopreservation method for *A. platensis* strain NIES-39 and adapt this method for an upcoming spaceflight experiment. Seven cryoprotective agents were tested to preserve viable *A. platensis*. We found that a combination of dimethyl sulfoxide and trehalose additives to Zarrouk’s media protected trichome viability for extended storage periods at -80 °C. Throughout one year of frozen storage, we observed equivalent viability of cryopreserved cells in comparison to non-frozen cultures. Equivalent cryopreservation was demonstrated with multiple volumes of frozen culture, culture containers, and thawing methods to adapt the method for delivery and return of viable NIES-39 for the Space Algae-2 experiment on the International Space Station. The method requires minimal resources, employs simple freezing and thawing procedures, and could be implemented in commercial production to preserve genetic stocks.

## 1. Introduction

*Arthrospira platensis* is an edible, filamentous cyanobacterium with a spiral trichome shape. Commonly known as spirulina, it is commercially cultured for pigments, nutritional supplements, biofuel, wastewater treatment, and CO_2_ sequestration^1^. Spirulina is also a promising organism for bioregenerative life support systems in long-duration space missions^2,3^. This cyanobacterium contains high levels of protein, antioxidants, and B-vitamins among other micronutrients^4^. Spirulina does not require preprocessing steps for human consumption, because the thin 40-60 nm cell wall is made up of 86% digestible polysaccharides^5^. Spirulina production in open bioreactor systems is more economical and commonly implemented for commercial production^6^. However, systems open to the natural environment can become contaminated with other microorganisms creating challenges for axenic production of *A. platensis*^7^. Axenic stocks of spirulina are maintained through serial subculturing, which requires regular culture passages. Subcultures carry risks of genetic drift within the stock cell population and contamination of the stock with other microorganisms, while lyophilization or cryopreservation of algae can be long-term storage solutions that preserve genetic integrity of algae stocks^8^.

Desiccation of spirulina causes significant loss of cell viability, metabolic changes, and a long reactivation phase for the low number of surviving cells^9^. The National Institute for Environmental Studies (NIES) collection developed cryopreservation methodology using 3% dimethyl sulfoxide (DMSO) as a cryoprotective agent (CPA) that preserved varying levels of viability for 136 of 144 cyanobacteria strains, although neither *A. platensis* strain of two tested showed efficient preservation^10^. A stepwise freezing approach using multiple CPAs and thawing at 40 °C resulted in 0-5% survival of two spirulina strains after seven months of cryopreservation^11^. A more robust *A. platensis* cryopreservation method was reported by Shiraishi^12^, in which multiple concentrations of DMSO and methanol were analyzed for seven *Arthrospira* strains. Cell survival was variable and dependent on strain genotype. For example, NIES-39 required stringent freezing and thawing procedures to maintain viability with preservation in 10% DMSO in Spirulina-

Ogawa-Terui (SOT) medium. After one year of storage at -80 °C there was ∼50% loss of viability, and the culture was stable with a 60% survival rate after storage in liquid nitrogen vapor phase^12^.

*A. platensis* strain NIES-39 is the subject of study for long-duration, crewed spaceflight as a component of bioregenerative life-support research and as a source of nutrition^13^. The planned Space Algae-2 experiment will test continuous subculturing of NIES-39 for 5-6 months on the International Space Station (ISS). The experiment requires a method for delivery of viable *A. platensis* to the ISS with flexible culture activation time and for return of preserved viable cells for genetic screening. It is possible to store viable spirulina cells without cryopreservation. Up to 10 weeks of storage was demonstrated for NIES-39 using heterotrophic dark conditions with Zarrouk’s medium (ZM), supplemented with glucose, in fluorinated ethylene propylene (FEP) bioreactor bags^13^. Similarly, a *Limnospira indica* spirulina strain can be stored for two weeks at 4 °C with low light levels^14^. These short-term storage methods can reduce the frequency of subculturing passages needed to maintain spirulina strains, but long-term cryopreservation is desirable to initiate cultures at any time in spaceflight and to return viable cells.

Here we report a long-term cryopreservation method for *A. platensis* NIES-39 suitable for spaceflight applications. Empirical tests of CPA combinations showed that 10% DMSO, 10% trehalose in ZM preserves full cell viability when frozen at -80 °C for one year. This method does not require specialized freezing or thawing conditions and may be applicable to the spirulina culture industry to preserve genetic stocks of individual strains without the risk of viability loss.

## 2. Methods

### 2.1. Strain and culture medium

*Arthrospira platensis* NIES-39 (UTEX 3086, Culture Collection of Algae, University of Texas at Austin) was used to test each cryopreservation method. ZM without vitamin B_12_ at pH 9 was used as the culture media^15^.

### 2.2. Cell preparation methods to test 10% DMSO cryopreservation

DMSO at 10% (v/v) was tested as a cryoprotectant with a modified method from Shiraishi^12^ that used ZM instead of SOT media. Alternative cell preparation methods were compared starting with a dense culture of approximately 1.6 optical density at 750 nm (OD_750_) as measured with either a GENESYS™ 180 UV-Vis Spectrophotometer (Thermo Fisher Scientific) or an M3 SpectraMax Microplate Reader (Molecular Devices). Figure 1 outlines the three methods for cell preparation, CPA addition, and cell recovery at stored volumes of 50 µL and 500 µL. The resuspension method centrifuged 500 µL of dense culture at 5,000 g for 5-8 min. Supernatant media was removed, cell pellets were resuspended in 500 µL of 10% DMSO in ZM, and 50 µL aliquots were cryopreserved. For the CPA concentrate method, 900 µL of dense culture was mixed with 100 µL DMSO for a final concentration of 10% DMSO and 50 µL samples were aliquoted for cryopreservation. Third, the 1:1 mix method combined 1 mL dense culture with 1 mL 20% DMSO in ZM for a final 10% DMSO concentration, and 500 µL samples were aliquoted. The samples were held at room temperature (RT) for at least 15 minutes and frozen to -80 °C at a rate of -1 °C min^-1^ in a Mr. Frosty^TM^ Freezing Container (Thermo Fisher Scientific). Samples were frozen for 24 h prior to thawing and assaying for viability. For the 1:1 mix method, frozen storage periods of 5 and 14-16 days were also tested.

**Figure 1.**
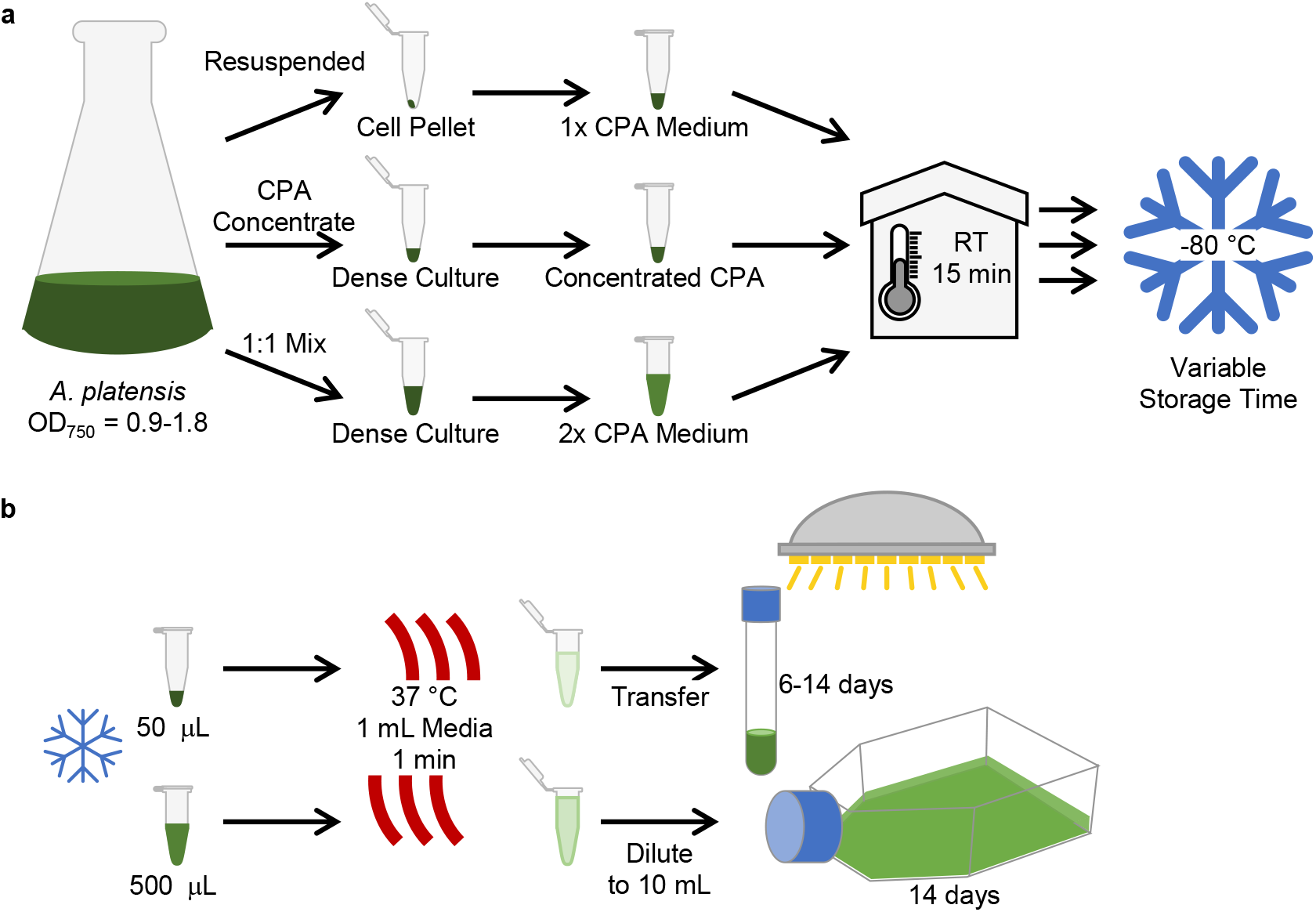
(a) Schematic of cryopreservation methods. *A. plantensis* cultures were grown to 0.9 to 1.8 OD_750_ in ZM. Three methods were used to mix cells and CPAs at desired concentrations. Resuspended cells were pelleted by centrifugation and resuspended in ZM media containing 1x concentration of the CPA. The CPA concentrate method directly added 10x or 20x CPA solution to the dense culture for a 1x final concentration. The 1:1 mix method combined equal volumes of culture with 2x CPA in ZM. Either 50 µL or 500 µL aliquots were cryopreserved, and cells were frozen at -80 °C for storage times up to one year. (b) Recovery of cryopreserved samples. Both 50 µL and 500 µL aliquots were thawed with 1 mL 37 °C ZM and the sample was held in a 37 °C water bath for 1 min. This diluted the 50 µL samples 1:20 of the cryopreserved volume prior to transfer to culture tubes. The larger volume samples of 1.5 mL after thawing were further diluted in ZM to a 1:20 dilution of cryopreserved volume (10 mL final volume) in tissue culture flasks. Tubes were incubated at 24 °C and flasks at RT with light to promote photosynthetic growth.

To assay for viability, sample tubes were removed from the -80 °C freezer in a Mr. Frosty^TM^ Freezing Container to keep frozen until thawed. For 50 µL samples, 1 mL of 37 °C ZM was added, and the tube was incubated in a 37 °C water bath for 1 minute. The thawed culture was transferred to a 10 mL round bottom tube and cultured for 6-7 days with 20 ± 4 µmol m^-2^ s^-1^ light at 24 °C in a Infors HT Multitron incubator. Light intensity was measured with an Apogee Instruments MQ-200 quantum light meter. The 500 µL samples were also mixed with 1 mL of 37 °C ZM and incubated in a 37 °C water bath for 1 minute. The samples were then transferred to 8.5 mL of 37 °C ZM in a 25 mL tissue culture flask with a vented cap. The flasks were cultured for 14 days at RT (22-26 °C) under white LED lights (Cat# C421510, Home Depot) at 20 ± 4 µmol m^-2^ s^-1^ light.

### 2.3. Sample analysis

Cryopreservation efficiency was determined by comparing the growth of a cryopreserved sample post thaw to a mock CPA, non-frozen, control or no CPA, non-treated cells. Mock CPA controls were prepared from the same inoculum of NIES-39 culture (OD_750_ = 0.9-1.8) used for each cryopreservation experiment. The mock CPA non-frozen controls followed the corresponding cell preparation method, i.e. resuspended, CPA concentrate, or 1:1 mix methods. Instead of freezing, three aliquots of these cells were diluted with ZM following the identical thawing procedures for each trial and grown for 14-17 days and growth was measured by absorbance at OD_750_ every two to three days. For no CPA cell controls, the same inoculum used for CPA treatment was directly diluted into ZM to collect absorbance measurements over 14-17 days. Cryopreserved aliquots were thawed after a frozen storage period and grown in identical conditions to the corresponding fresh cell control with absorbance measurements at OD_750_ every two to three days. For each inoculum preparation, OD_750_ measurements were compared for the same day of growth during mid-log phase for both the cryopreserved and fresh cell treatments. These paired absorbance measurements were used to calculate the percent recovery of viable cells as [OD_750_(Frozen)/OD_750_(Fresh)] x 100%. Specific growth conditions are described below for each experiment.

### 2.4. Cryoprotectant screen

Including 10% DMSO, eight CPA additions to ZM were tested for cryoprotective properties. Final concentrations for each CPA tested in ZM were: 10% glycerol, 5% methanol, 10% methanol, 5% glycine betaine, 10% trehalose, 10% DMSO with 10% Ficoll-70, 10% DMSO with 0.1% polyethylene glycol (PEG) 8000, and 10% DMSO with 10% trehalose. Percentages are given w/v for solids and v/v for liquids at RT. A slow freeze of -1 °C min^-1^ was used for tests except for the following samples, which were frozen in a -80 °C freezer with a cardboard cryobox: 10% trehalose samples, nine 1-day 10% DMSO samples, and five 14-to 16-day 10% DMSO samples. All samples were thawed and diluted 1:20 with 37 °C ZM except for 10% glycerol samples, for which 300 μL of the 1.5 mL thawed samples was inoculated into 10 mL cultures. Post thaw, the cells were grown either in a Multitron incubator at 24 °C or under white LED lights at RT (22-26 °C) with 20 ± 4 µmol m^-2^ s^-1^ illumination.

### 2.5. Long-term cryopreservation

Long-term stability was tested with NIES-39 preserved in 10% DMSO with 10% trehalose in ZM for 365 days. The cryoprotected NIES-39 culture was prepared with the 1:1 mix method with 500 µL aliquots. Three samples were processed for the non-frozen control recovery assay, and the remaining aliquots were frozen in a cardboard cryobox at -80 °C. At each storage timepoint, three samples were thawed following the 1:1 mix method and diluted to 10 mL cultures in tissue culture flasks. The cultures were held at RT for 30 min with ambient laboratory lighting prior to 14 days at 24 °C with 20 ± 4 µmol m^-2^ s^-1^ continuous light in a Multitron incubator. Absorbance measurements for each recovery assay were fitted to a logistic growth curve using the R package, Growthcurver^16^. The Growthcurver t-gen metric estimates doubling time.

At the 264-day timepoint, the 1:1 mix thawing method at 37 °C was compared to an additional three samples that were thawed at RT and diluted to 10 mL culture volume with ZM at RT. One hour post thaw, RT and 37 °C thawed samples were imaged on a Nikon Diaphot inverted microscope at 40X magnification with an AmScope 5 MP High-Speed Color CMOS C-Mount Microscope Camera.

### 2.6. Stability of CPA stock solution

ZM with 20% DMSO and 20% trehalose was stored at RT, protected from light, in a 50 mL conical tube for 7.5 months. A dense *A. platensis* culture was used to test biocompatibility and effectiveness of the aged CPA solution. For a fresh inoculum, no CPA control, three replicate 10 mL cultures were inoculated directly with 500 µL of the dense culture. To test cryopreservation, the stored CPA solution was 1:1 mixed with six 500 µL replicate samples from a dense *A. platensis* culture. Three samples were diluted to 10 mL culture volume without freezing and incubated at 24 °C with 20 ± 4 µmol m^-2^ s^-1^ continuous light for 15 days in a Multitron incubator. The final three samples were cryopreserved and kept frozen at -80 °C for 15 days. After 15 days of frozen storage, the cryopreserved samples were thawed, mixed with 1 mL 37 °C ZM, held 1 min at 37 °C, and then diluted to 10 mL culture volume for 15 days of culture in a Multitron incubator. Growth was assayed by OD_750_ for all cultures.

### 2.7. Adaptation for spaceflight

Feasibility of cryopreserved culture delivery to the ISS was tested using three different concepts of operations. First, *A. platensis* dense culture was 1:1 mixed with 20% trehalose, 20% DMSO in ZM. A 2 mL volume of the 1:1 mix was added to American Durafilm, 14.5 cm × 15.2 cm, FEP culture bags (0.127 mm, 5 mil thickness). Three no CPA non-frozen control culture bags were initiated by diluting 1 mL of the identical dense culture for CPA treatment with 99 mL of 37 °C ZM. The non-treated controls were immediately cultured for 15 days in a Multitron incubator at 24 °C with 20 ± 4 µmol m^-2^ s^-1^ light. Cryopreserved FEP bags were frozen at -80 °C inside metalized food bags (Uline, CAT # S-18133) with a zipper-top closure and stored for 14 days. To thaw, six frozen bags were exposed to RT for approximately 5 minutes and then 37 °C ZM (3 bags) or RT ZM (3 bags) was added to a final volume of 100 mL. The FEP culture bags were incubated for 15 days in a Multitron incubator at 24 °C with 20 ± 4 µmol m^-2^ s^-1^ light.

Second, 2 mL aliquots of CPA-treated *A. platensis* cells in 10% DMSO and 10% trehalose in ZM were transferred into 3 mL syringes, sealed with Breathe-Easy® membrane (Diversified Biotech), and frozen at -80 °C. After 21 days frozen storage, three syringes were thawed at 37 °C and mixed with 98 mL ZM in FEP culture bags. Three no CPA control cultures were inoculated from the original dense culture on the day of cryopreservation. All cultures were grown in a Multitron incubator at 24 °C with 20 ± 4 µmol m^-2^ s^-1^ light.

Third, trehalose without DMSO was tested as a CPA. Briefly, 2 mL of 1:1 mix cells containing 10% trehalose in ZM were cryopreserved in FEP bags and frozen for 33, 55, or 69 days. Three no CPA control culture bags were cultured at the zero time point of the cell preparation and initial freeze. Cryopreserved samples on day 33 and 69 were thawed and diluted at RT. The 55-day samples were thawed and diluted at 37 °C. All FEP bag cultures were grown at 24 °C with 20 ± 4 µmol m^-2^ s^-1^ light in a Multitron incubator for 17 days.

## 3. Results

We initially tested a modified cryopreservation sample preparation method from Shiraishi^12^ in which dense cultures of *A. platensis* NIES-39 were centrifuged to pellet the cells and then resuspended in ZM with 10% DMSO. After 24 h frozen storage, this resuspended method resulted in an average recovery of 48% the growth of non-frozen controls with a standard deviation (SD) of 26% (N= 8). To simplify the methodology for spaceflight applications, we tested mixing equal volumes of 20% DMSO in ZM with dense culture prior to the 24 h freeze. This 1:1 mix had 52 ± 29% (N = 13) viability after cryopreservation compared to non-frozen, mock CPA, control cultures. The resuspended and 1:1 mix methods were not statistically different based on a two-tailed Student’s t-test (p = 0.77). We also tested a CPA concentrate addition method in which DMSO was added directly to the dense culture to a final concentration of 10% DMSO. The direct addition of DMSO produced no viable cells after thawing (N = 4). We then focused on the resuspended and 1:1 mix methods for longer frozen storage time periods. Figure 2 shows loss of viability with relatively short frozen storage periods of five and 14 days with both cell preparation methods indicating that 10% DMSO is not an optimal CPA for the NIES-39 strain.

**Figure 2.**
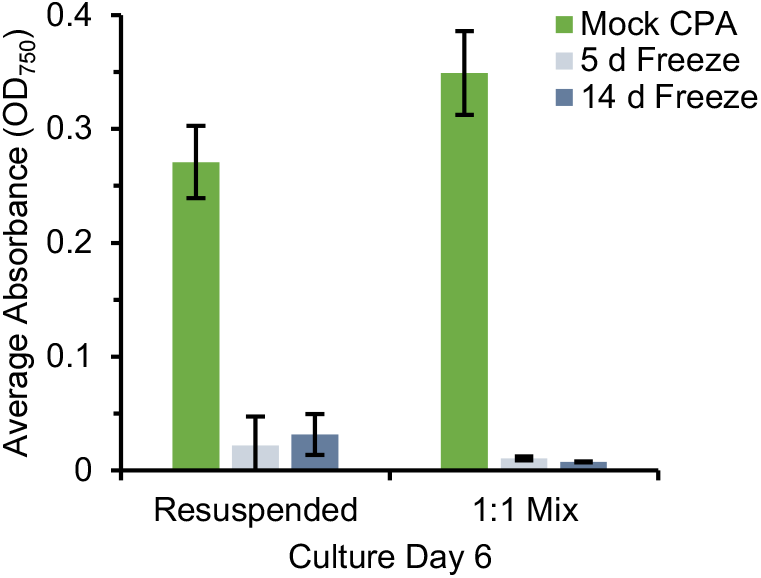
Effect of frozen storage time on NIES-39 viability with 10% DMSO as the CPA. The mock CPA treatment cultured non-frozen cell aliquots from the 10% DMSO treatments with a mock thawing procedure. Frozen samples were thawed with 37 °C medium, and cell viability was assayed by optical absorbance at 750 nm after incubating with light for 6 days. Average and SD of three replicates per treatment are plotted. The low absorbance values for the frozen cell methods are equivalent to typical values of the thawed, diluted cryopreserved inoculum at initiation of the recovery growth assay.

To identify CPAs that would better support long-term frozen storage of NIES-39, we tested glycerol, methanol, glycine betaine, and trehalose as CPAs. In addition, we tested Ficoll-70, PEG 8000, and trehalose in combination with DMSO. Table 1 summarizes the cell preparation and cryopreservation conditions tested as well as recovery descriptive statistics from these trials. Both freshly prepared mock CPA control and frozen cultures for the 10% glycerol CPA treatment were not viable. Methanol at either 5% or 10% in ZM had low viability with only 3-4% recovery from a 24 h freeze. Some delayed regrowth, after 17 days of incubation, was observed in one 5% methanol and two 10% methanol recovery cultures. The 5% glycine betaine CPA treatment was frozen for 16 days and resulted in 10% recovery. By contrast, 10% trehalose in ZM had an average recovery of 76% after freezing for 17 days indicating that trehalose was more cryoprotective than DMSO for long-term storage.

**Table 1.** Summary of CPA tests with *A. platensis*. Percent recovery is shown for 6-7 days post thaw with illumination of 20 ± 4 µmol m^-2^ s^-1^ compared to non-frozen mock CPA controls cultured at the time of initial CPA treatment.

| Final Conc. of CPA in ZM | Input OD<sub>750</sub> | CPA addition | Vol. Frozen (μL) | Days Frozen | Recovery Lighting | Sample Size | Mean Recovery ± S.D. (%) |
| --- | --- | --- | --- | --- | --- | --- | --- |
| 10% DMSO | 1.6-1.8 | 1:1 Mix | 500 | 1 | Multitron | 10 | 50 ± 35 |
| 10% DMSO | 1.4 | 1:1 Mix | 500 | 1 | LED | 3 | 59 ± 4 |
| 10% DMSO | 1.1-1.7 | 1:1 Mix | 500 | 14-16 | LED | 8 | 3 ± 1 |
| 10% Glycerol | 1.1 | 1:1 Mix | 500 | 1 | Multitron | 6 | Not Viable |
| 10% Methanol | 0.9 | Resuspend | 50 | 1 | LED | 2 | 3 ± 0 |
| 5% Methanol | 0.9 | Resuspend | 50 | 1 | LED | 2 | 4 ± 0 |
| 5% Glycine Betaine | 1.4 | 1:1 Mix | 50 | 16 | Multitron | 3 | 10 ± 2 |
| 10% Trehalose | 1.6 | 1:1 Mix | 500 | 17 | Multitron | 5 | 76 ± 5 |
| 10% DMSO, 10% Ficoll-70 | 1.7 | 1:1 Mix | 500 | 14-16 | LED | 5 | 25 ± 19 |
| 10% DMSO, 0.1% PEG 8000 | 1.5 | 1:1 Mix | 50 | 14 | Multitron | 3 | 58 ± 14 |
| 10% DMSO, 10% Trehalose | 1.7 | 1:1 Mix | 500 | 16 | LED | 2 | 102 ± 14 |

We also tested three CPAs in combination with 10% DMSO (Table 1). Compared to 10% DMSO alone, all three additives improved viable cell recovery after 14-16 days of frozen storage. Addition of 10% Ficoll-70 improved recovery from 3% for DMSO alone to 25%, while addition of 0.1% PEG 8000 increased recovery of viable cells to 58% of non-frozen, mock CPA control cultures. The combination of 10% DMSO and 10% trehalose in ZM resulted in 102% of growth compared to mock CPA controls, which suggested little to no viability loss with a combination of DMSO and trehalose as the CPA.

Cryopreservation with 10% DMSO, 10% trehalose in ZM was tested at nine timepoints over 365 days of frozen storage. Figure 3a shows the growth of three non-frozen, mock CPA aliquots over 14 days at the start of the storage experiment. Figure 3b-d show equivalent recovery growth assays for three replicates each that were thawed on 105, 264, and 365 days of frozen storage. Logistic growth curves were fitted to the individual recovery assays using the Growthcurver R package^16^, which estimated doubling time for cell recovery at each storage timepoint based on non-linear least-squares Levenberg-Marquardt fits (Figure 3e). The doubling times did not show statistically significant differences based on storage time (p = 0.16, single factor ANOVA). Percent recovery of each sample was calculated by comparing the 9-d OD_750_ of the individual samples to the average 9-d OD_750_ for the mock CPA controls (Figure 3f). A single factor ANOVA showed significant differences based on storage time (p = 2 ×10^-8^). Tukey’s HSD tests identified storage days 20 and 226 as being significantly lower and higher percent recovery, respectively, compared to all other assays. Both the doubling time estimates and percent recovery assays suggest that there is no loss in *A. platensis* NIES-39 viability with up to a year of -80 °C frozen storage in 10% DMSO and 10% trehalose in ZM.

**Figure 3.**
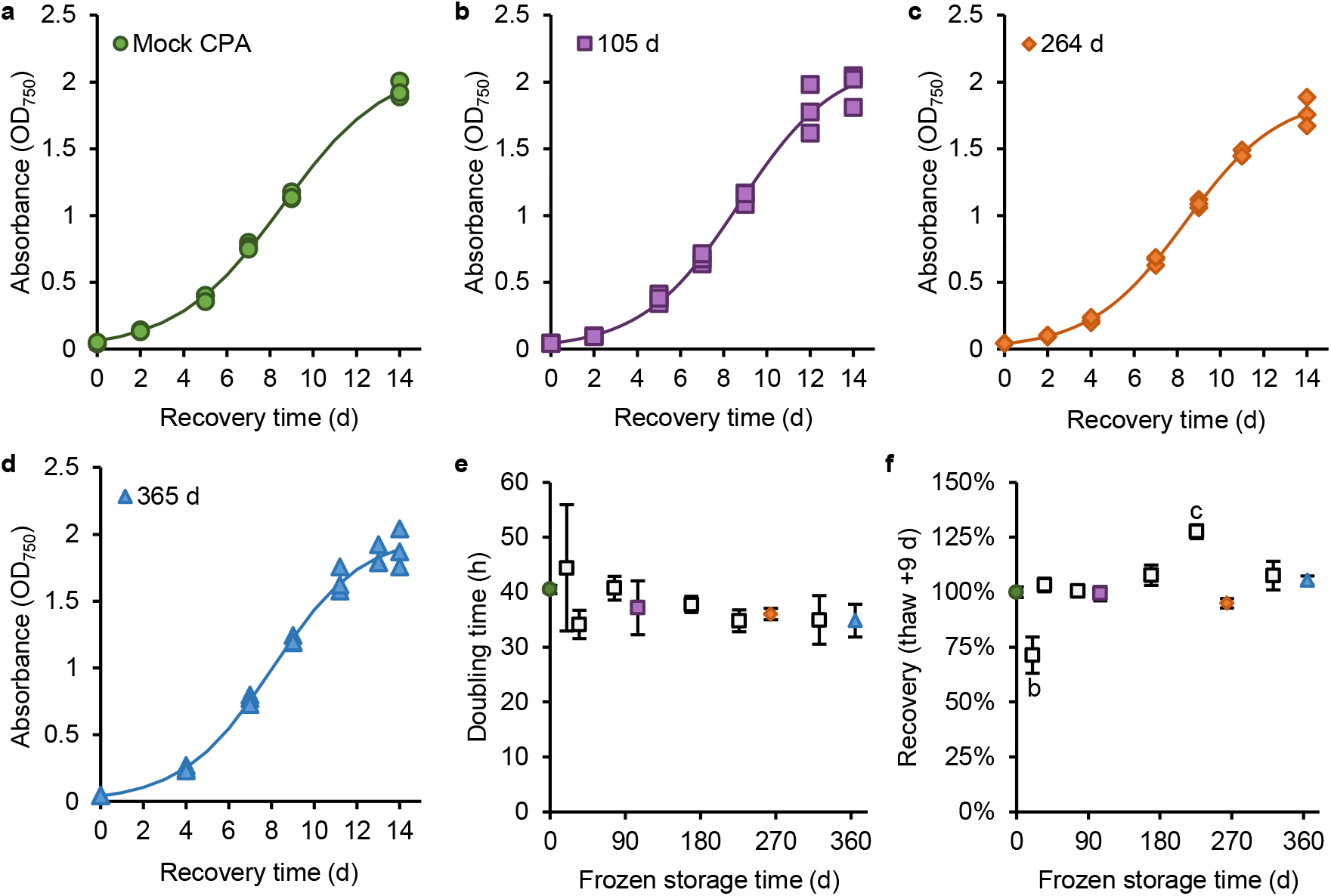
Effect of frozen storage time on cryopreservation of *A. platensis* cultures in ZM with 10% DMSO and 10% trehalose. (a-d) Example recovery assays from: non-frozen, mock CPA control (a), 105 days frozen storage (b), 264 days frozen storage (c), and 365 days frozen storage (d). Three replicate samples were assayed at each timepoint. Lines are logistic growth curves fit to the combination of the three replicate assays. (e) Average doubling time estimated from logistic growth curve fits for each recovery assay over ten storage timepoints. Error bars are SD. Single factor ANOVA based on storage time was not significant (p = 0.16). (f) Average recovery percentage at day 9 of the recovery assay for each storage timepoint. Error bars are SD. Letters b and c indicate storage timepoints that were significantly different from all other timepoints based on a single factor ANOVA (p = 2 ×10^-8^) and post-hoc Tukey’s HSD tests (p < 0.01 for all significant tests, p > 0.1 for all non-significant tests). (e-f) The colored symbols are derived from the data shown in a-d as follows: the green circle is the mock CPA control; the purple square is 105 days storage; the blue diamond is 264 days storage; and the red triangle is 365 days storage.

We then tested a series of cryopreservation and recovery method modifications to simplify implementation of cryopreservation in spaceflight. A thaw and cell recovery with media at 37 °C and a 1 min 37 °C incubation prior to the growth recovery assay requires equipment, crew engagement, and time for warming the medium and incubator. We tested whether thawing cryopreserved cells with RT medium impacted the 264-d long-term frozen storage timepoint. In addition to the three samples thawed at 37 °C, three frozen samples were thawed by adding 1 mL ZM at RT. Predominantly intact trichomes were observed 1 h after thawing at both 37 °C and RT (Figure 4a-b). Figure 4c compares recovery growth curves of thawed samples at both temperatures. The 9-d percent recovery was 95 ± 2% and 88 ± 2% for the 37 °C and RT thawed conditions, respectively. Although the 9-d recovery was significantly lower in the RT thaw (Student’s t-test, p = 0.3), the recovery growth curves are not significantly different based on two factor ANOVA (p = 0.13 for the thawing condition effect).

**Figure 4.**
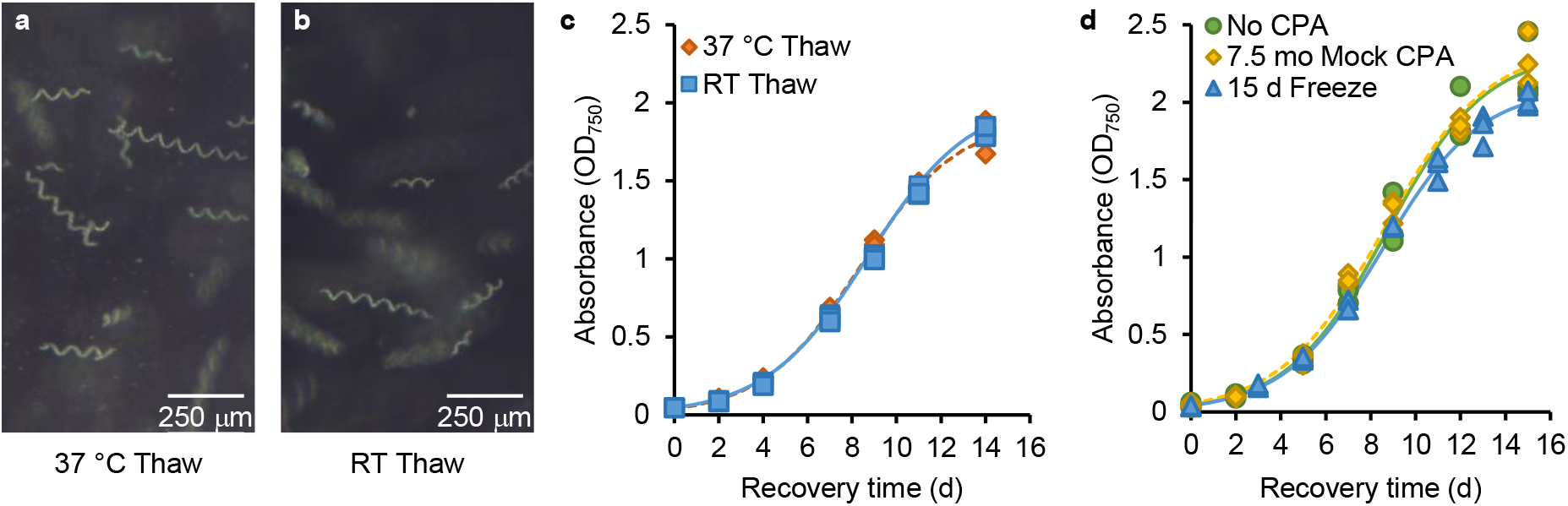
Cryopreservation modifications to improve spaceflight implementation of the 10% DMSO with 10% trehalose CPA. (a-b) Darkfield micrographs of *A. platensis* cultures thawed either at 37 °C (a) or at RT (b) after 264 days of frozen storage. The trichomes were imaged 1 hour after thawing. (c) Recovery growth curves comparing the 37 °C (orange diamonds) and RT (blue squares) thawing treatments for the 264-d storage timepoint. The solid blue and dotted orange lines are logistic growth curve fits of three samples per treatment using the Growthcurver R package. (d) Cryopreservation effectiveness of 20% DMSO with 20% trehalose in ZM after RT, dark storage for 7.5 months. No CPA, non-frozen control cells (green circles) were compared with non-frozen cells treated with the CPA solution (yellow diamonds), and CPA-treated cells frozen for 15 days (blue triangles). Corresponding green, yellow, and blue lines are logistic growth curve fits of three samples per treatment.

To preserve cells grown in space, pre-mixed 20% DMSO with 20% trehalose in ZM will need to be supplied from Earth and will likely be stored at RT in the dark. We tested the stability and biocompatibility of the CPA solution after 7.5 months of RT, dark storage. Figure 4d compares growth of non-frozen cells without CPA (No CPA), non-frozen cells mixed with the 7.5-month-old CPA solution (7.5 mo Mock CPA), and cryopreserved cells held at -80 °C for 15 days. At 9-days of growth, the average percent recovery of the mock CPA cells was 107 ± 6%, while the average percent recovery of the 15-day frozen cells was 98 ± 0.3%. Single factor ANOVA of percent recovery, including the no CPA control cells, was not significant (p = 0.47) indicating that the stored CPA had no impact on cell viability.

The cryopreservation method was then adapted for delivery of viable cells to the ISS. The cryopreserved volume was scaled to 1 mL of dense culture (OD_750_ = 2.03) mixed with 1 mL of 20% DMSO 20% trehalose in ZM, which was frozen in an FEP tissue culture bag to provide sufficient inoculum for a 100 mL culture Figure 5a. Three no CPA, non-frozen control cultures were initiated with 1 mL of the same *A. platensis* inoculum into 100 mL total culture volumes in FEP bags. Frozen samples were thawed after 14 days of storage with either 37 °C or RT ZM. Average percent recovery at 9 days of post-treatment culture was 98 ± 14% for the 37 °C thaw and 105 ± 7% for the RT thaw. A single factor ANOVA for percent recovery was not significant (p = 0.66). Figure 5b plots the absorbance data and fitted logistic growth curves showing extensive overlap between the three treatments. These data suggest that the cryopreservation method is scalable and biocompatible with FEP tissue culture bags.

**Figure 5.**
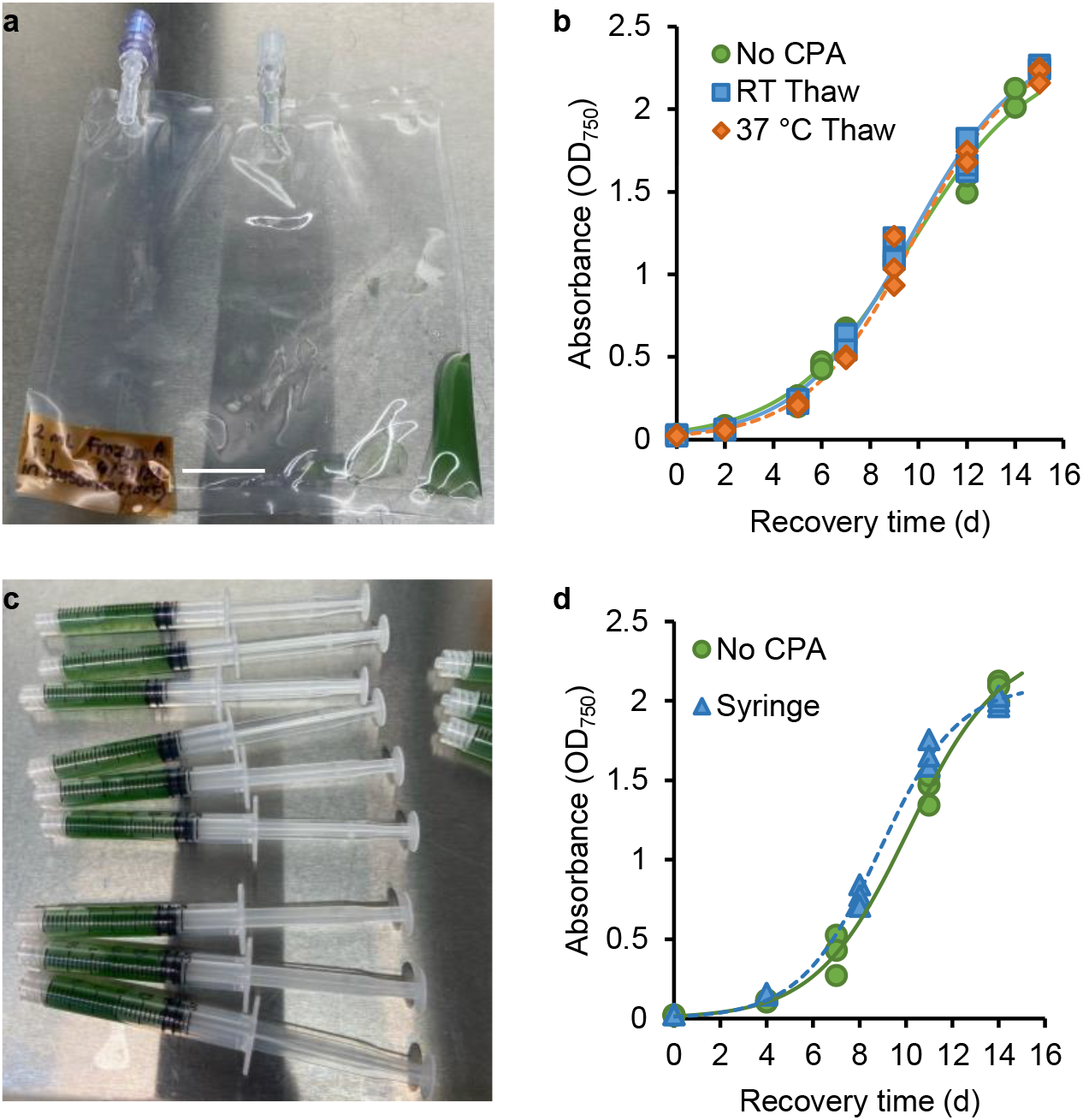
Adaptations of cryopreservation to spaceflight-like hardware. (a) CPA-treated cells in an FEP culture bag prior to cryopreservation, (b) Recovery of cells after 14 days of frozen storage. Frozen FEP bags were thawed with 98 mL of 37 °C (orange diamonds) or RT (blue squares) ZM. The no CPA treatment (green circles) was a 1 mL inoculation of the dense used for cryopreservation into an FEP bag with 99 mL of ZM. Lines are logistic growth curve fits of three replicates per treatment. (c) Preparation of 3 mL syringes with 1:1 Mix *A. platensis* cells with 10% DMSO, 10% trehalose in ZM. (d) Comparison of growth of non-treated cells (green circles) and cryopreserved (blue triangles) samples after 21 days of frozen storage in syringes. Lines are logistic growth curve fits of three replicates per treatment.

Freezer volumes are limiting in spacecraft. To reduce the container volume of frozen inoculum, we tested whether 3 mL syringes could be used for cryopreservation of *A. platensis* (Figure 5c). Three syringes were thawed at 37 °C after 21 days of frozen storage and used to inoculate 100 mL cultures in FEP bags. Figure 5d plots growth of no CPA, non-frozen control cells inoculated from the original dense culture on the day of cryopreservation versus frozen cells stored for 21 days in syringes. Recovery of the frozen cells at 11 days of culture was 115 ± 6% relative to the non-treated control cells.

It may be desirable to eliminate DMSO, an organic solvent, from *A. platensis* cryopreservation in spacecraft. Trehalose without DMSO was tested as the CPA for frozen delivery of *A. platensis* to the ISS. *A. platensis* was cryopreserved in an FEP bag with 10% trehalose in ZM. After freezing at -80 °C, the cyanobacteria were thawed with RT ZM on day 33 and 69 of storage and thawed with 37 °C ZM on day 55 of storage. Figure 6a plots recovery growth and logistic growth curve fits for no CPA non-frozen control cells as well as the frozen storage timepoints with different thawing temperatures. Figure 6b plots percent recovery growth at 8 days of culture, which was significantly reduced for both RT thaw trials (single factor ANOVA p = 1.15 x 10^-5^). Tukey HSD tests indicated the non-treated control cells and 37 °C were not significantly different from each other (p = 0.35) and that both RT thaw trials were not significantly different (p = 0.90). However, the RT thaw trials were significantly different from the control and 37 °C thaw trials (p < 0.01 for all comparisons). These data suggest that 10% trehalose alone as the CPA increased the sensitivity of the frozen cells to the thawing temperature.

**Figure 6.**
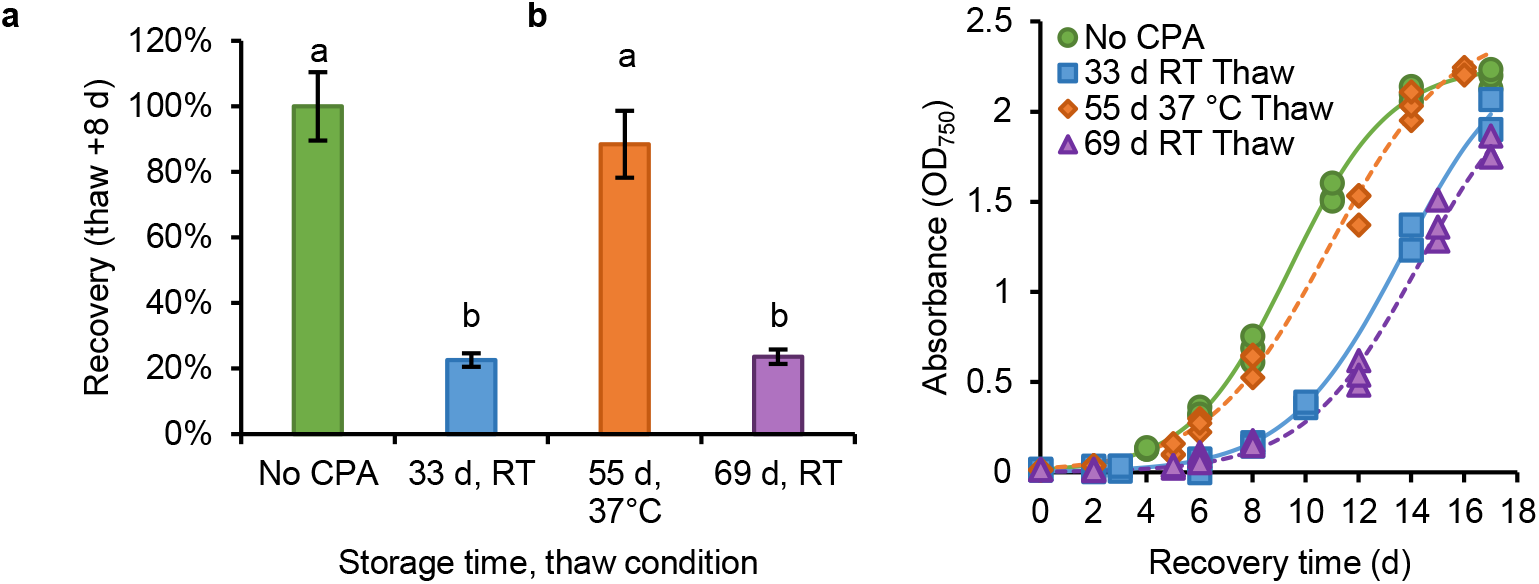
Comparison of recovery growth with different frozen storage time and thawing temperature conditions when 10% trehalose was the sole CPA in FEP tissue culture bags. (a) FEP bags were thawed with 98 mL of 37 °C after 55 days of frozen storage (orange diamonds) or with RT ZM after 33 (blue squares) or 69 days (purple triangles) of frozen storage. The no CPA treatment (green circles) was a 1 mL inoculation of the dense used for cryopreservation. Lines are logistic growth curve fits of three replicates per treatment, except for two replicates at the 33-day RT thaw treatment. (b) Average recovery percentage at day 8 of the recovery assay for each storage and thaw treatment. Error bars are SD. Letters a and b indicate treatments that are significantly different based on a single factor ANOVA (p = 1.5 ×10^-5^) and post-hoc Tukey’s HSD tests (p < 0.01 for all significant tests, p > 0.35 for all non-significant tests).

## 4. Discussion

Prior reports of *A. platensis* cryopreservation suggest that most strains are highly sensitive to frozen storage^10–12^. The CPA, freezing method, storage temperature, and thawing conditions all influence *A. platensis* freeze tolerance. Shiraishi^12^ showed that 10% DMSO, slow freezing, storage in liquid nitrogen vapor, and rapid warm temperature thawing were the most optimal of the conditions tested for maintaining viable *A. platensis* NIES-39 in long-term frozen storage. The Space Algae-2 ISS experiment is designed to grow *A. platensis* NIES-39 in spaceflight, but storage in liquid nitrogen vapor is not feasible for launch and transport to the ISS. We initially tested whether modified cyanobacteria growth, freezing, and thawing methods using 10% DMSO as the CPA would provide sufficient cryoprotection. However, we found a rapid decline in viability with no recovery of viable cells after a 5-d storage period at -80 °C suggesting that 10% DMSO is not a robust CPA for NIES-39 trichomes when grown in culture conditions that deviate from the published method^12^.

DMSO is used widely as a CPA due to its properties of penetrating cell membranes and inhibiting damaging ice nucleation during freezing and thawing of cells^17,18^. Although DMSO is more effective at inhibiting ice nucleation at higher concentrations, it also becomes more toxic to cells as concentration increases, which limits the amount of DMSO that can be added as a CPA. Exogenous sugars, such as trehalose, and other metabolites, such as glycine betaine, can also be effective CPAs and are less toxic to cells. These metabolites can act as both extracellular and intracellular CPAs but are not membrane permeable and require membrane protein transporters to provide intracellular ice protection^18^. Exogenous glycine betaine mitigates high-salt stress growth response in *A. platensis* suggesting potential to mitigate other abiotic stresses^19^. We observed a limited cryoprotective effect of glycine betaine with significantly delayed growth (Table 1). Despite glycerol being a common CPA, mixotrophic culture of *A. platensis* with glycerol can be inhibitory of growth^20,21^. Even without freezing, we observed no growth after treatment with 10% glycerol.

By contrast, there is more evidence that trehalose was expected to be a robust CPA. *A. platensis* produces trehalose endogenously in response to high salt environmental stress suggesting that trehalose accumulation is part of wild-type environmental stress adaptation^22^. Exogenous trehalose can mitigate low nitrogen stress in *A. platensis* and promote glycogen as well as poly-3-hydroxybutyrate accumulation suggesting this cyanobacterium may uptake trehalose^23^. We found that 10% trehalose alone was a more effective long-term CPA compared to 10% DMSO alone. After 55 days of frozen storage, we observed 88 ± 10% recovery when frozen trichomes were rapidly thawed at 37 °C (Figure 5d). However, trehalose as a sole CPA may require controlled thawing conditions to maintain cell viability, because thawing in RT medium at 33 or 69 days of frozen storage resulted in much lower recovery of viable cells.

The combination of 10% DMSO with 10% trehalose in ZM provided effective and adaptable cryopreservation. In nearly all experiments, this combination CPA showed equivalent growth between untreated, non-frozen, and cryoprotected trichomes using multiple storage containers including microcentrifuge tubes, FEP culture bags, and syringes as well as for frozen storage periods up to 365 days. Fast thawing at 37 °C was statistically equivalent to RT thaws at 14 and 264 days of frozen storage. Potentially, increasing the viscosity of the CPA solution by combining DMSO with trehalose reduces ice crystal nucleation during freezing, storage, and thawing^17^. It is also possible that the combination of cellular uptake of trehalose and permeation of DMSO promotes robust intracellular vitrification upon freezing, which would provide greater cryoprotective activity.

We also found that a 20% DMSO and 20% trehalose CPA in ZM can be stored for at least 7.5 months at RT and provide 14 days of cryoprotection for *A. platensis* relative to no CPA treatment. Long shelf-life of the CPA solution simplifies the cryopreservation method for implementation in spaceflight. In the Space Algae-2 experiment, space-grown cyanobacteria will be preserved up to 6-months after a serial batch culture experiment is initiated. After return to Earth, cryopreserved samples will be cultivated from individual trichomes to determine rates of genetic drift in space and ground control samples. The long shelf-life of the CPA solution allows for preservation of nearly all trichomes within a culture sample.

Demonstration of multiple variations in cryopreservation methods allow flexible experiment design and concepts of operations for launch and on-orbit cryopreservation. Compared to many biological experiments that are metabolically active during launch and transit to the ISS, cryopreserved cells allow for flexible timing of experiment initiation. By contrast, the Space Alage-1 experiment launched *Chlamydomonas reinhardtii* growing heterotrophically and required prompt activation of the experiment in a lighted growth chamber^24^.

The long-term viability of cryopreserved *A. platensis* reduces risk for incorporating edible cyanobacteria into bioregenerative life support systems. In the event of contamination of a spirulina production system in space, a cryopreserved stock could be used to restart production after a decontamination procedure. Without cryopreserved stocks, a separate spirulina stock culture would need to be continuously cultivated in addition to a biological production system. Likewise, this simple and effective cryopreservation method may have a positive impact on the spirulina cultivation industry and culture repositories to preserve stock cultures with reduced risk of accumulating mutations or loss of a genetic stock. The DMSO and trehalose CPA method does not require liquid nitrogen and is accessible to regions where only -80 °C freezers are present.

However, this study is limited to a single strain, NIES-39. In the future, both trehalose alone as well as DMSO with trehalose should be tested as CPAs for other *Arthrospira* species and strains.

## Acknowledgments

We thank Jared Brodderick for insightful discussions of our pre-publication data for this study.

